# Analysis of actin and focal adhesion organisation in U2OS cells on polymer nanostructures

**DOI:** 10.1101/2020.09.14.289330

**Authors:** Jakob Vinje, Noemi Antonella Guadagno, Cinzia Progida, Pawel Sikorski

## Abstract

Cells in their natural environment are embedded in a complex surrounding consisting of biochemical and biomechanical cues directing cell properties and cell behaviour. Nonetheless, *in vitro* cell studies are typically performed on flat surfaces, with clear differences from the more complex situation cells experience *in vivo*. To increase the physiological relevance of these studies, a number of advanced cellular substrates for *in vitro* studies have been applied. One of these approaches include flat surfaces decorated with vertically aligned nanostructures.

In this work, we explore how U2OS cells are affected by arrays of polymer nanopillars fabricated on flat glass surfaces. We focus on describing changes to the organisation of the actin cytoskeleton and in the location, number and shape of focal adhesions. From our findings we identify that the cells can be categorised into different regimes based on their spreading and adhesion behaviour on nanopillars. A quantitative analysis suggests that cells seeded on dense nanopillar arrays are suspended on top of the pillars with focal adhesions forming closer to the cell periphery compared to flat surfaces or sparse pillar arrays. This change is analogous to similar responses for cells seeded on soft substrates.

Overall, we show that the combination of high throughput nanofabrication, advanced optical microscopy, molecular biology tools to visualise cellular processes and data analysis can be used to investigate how cells interact with nanostructured surfaces and will in the future help to create culture substrates that induce particular cell function.

## 1 Introduction

*In vivo*, cells typically reside in a a complex 3D environment called extracellular matrix (ECM). The ECM not only serves as a structural scaffold for the cells, it is also a conveyor of biomechanical and biochemical signals and thus regulates a range of processes such as tissue morphogenesis, homeostatis and differentiation. It is composed of water, polysaccharides and proteins[1, 2, 3, 4], and the composition varies between tissue types.

Motivated by the need of creating cell culturing models that better represent *in vivo* conditions, researchers have increasingly started to study cell behaviour also in 3D matrices and in “semi-3D” systems. A number of differences in cell phenotypes between flat substrates and systems with higher dimensionallity have been identified [5, 6]. For example, characteristics such as viability, proliferation, differentiation and morphology are known to differ between cells on flat surfaces and cells embedded in 3D matricies[7, 3].

*In vivo*-like substrates range from “semi-3D”/2.5D substrates, such as flat surfaces decorated with various nanostructures to “true-3D” systems such as collagen gels or matrigel matrices[8, 9, 10, 11]. In addition, controlled positioning of ligands on surfaces may give new insights into how cells interact with various chemical patterns[12, 13, 14]. Also mechanical factors such as structure stiffness or even surface chemistry have been shown to influence cellular function[15, 16, 17]. To this end, a large number of different substrates for cellular studies have been developed[3, 18, 19, 20, 21, 22].

It has also been suggested that 3D culturing systems more precisely could predict the *in vivo* effect of a drug and thus these systems could find applications in drug discovery[16, 23, 24]. Precisely controlling nanoscale topographical patterns can also be used to regulate cell morphology. For example, wrinkles and grooves can be used to recreate the striated alignment of cardiomyocytes and thus better represent physiologically relevant conditions to model various diseases[25, 26].

The cytoskeleton of the cell is connected to the ECM is facilitated by focal adhesions (FAs), a multiprotein complex including cell surface integrins and scaffold proteins. Depending on a complex set of regulatory mechanisms, the FAs form and disassemble at a turnover rate needed for forward movement, for example in cell migration. The FAs are known to exert mechanical force on the ECM, and conversely the ECM exerting force on the cells is known to influence integrin affinity and avidity in the membrane[27].

One of the proteins known to be an integral part of the FAs is vinculin. It is one of the linker proteins involved in anchoring F-Actin to the integrin-complex. Lack of vinculin alters cell morphology, adhesion and motility[28], and impairs the cells ability to transduce force to the substrate[29, 30, 31]. Vinculin is not only involved in the mechanical connection of the actin cytoskeleton to the integrin-complexes, it also has the ability to crosslink and bundle actin filaments[32, 33, 34], modify existing actin bundles[35], cap actin filaments, nucleate new actin polymerisation sites[36] and recruit actin modifiers[37].

Cells respond to the 3D matrices by changing the number and type of cell-substrate adhesion and induce changes in the spatial organisation of the cytoskeleton. These changes in turn influence distribution, size and dynamics of the formed adhesions[38, 4, 39, 40, 41]. This rearrangement may lead to changes in cell proliferation, morphology and motility[42].

In order to understand the influence of complex 3D environments on cells, there is a need to develop new model systems where cellular processes can be studied and compared to flat controls. As cellular response is known to depend on physical, mechanical and chemical characteristics of the culturing substrate, it is desirable to fabricate cellular substrates with precisely controlled properties[43, 44, 45]. Additionally, it is highly advantageous if the cells and the substrate easily can be studied using already established analysis techniques such as optical microscopy.

One type of substrate that has recently gained attention are flat surfaces decorated with nanopillars or nano-wires[46, 47, 48, 49, 18, 50, 51, 52, 21, 53]. Compared to for example hydrogels, these structured surfaces do not mimic the true 3D environment, but have well defined surface topography. These substrates are typically referred to as being 2.5D. Such systems have already been applied to facilitate delivery of biologically relevant molecules into cells[54, 55], to monitor enzymatic activity[56], to test nuclear mechanics[57] and to study how tuning the membrane curvature influence various cell-membrane related processes[58, 59, 60]. By fabricating nanostructures on transparent substrates, it is possible to integrate this approach with optical microscopy.

The number possible combinations of different cell lines, nanostructure type and geometry is high, and examples from the literature are abundant. Li *et al.* described cell behaviour on surfaces decorated with randomly positioned gallium phosphide nanostructures and quantified the fraction of cells with large FAs[61]. The cell and FA morphology was investigated on surface with various area densities of nanowires. The results indicated that cells seeded on low-density surfaces were in contact with the substrate and formed large FAs around the cell edges. Large FAs were detected in a high fraction of cell on these arrays. For high nanowire areal densities and, cells were suspended on the top of the nanowire arrays and point-like FAs under the cells were observed. A lower fraction of cells on these arrays showed large FAs compared to cells on surface with low nanowire area density.

Buch-Månson *et al.* studied cell-nanostructured surface interactions for silicon nanocolumn arrays randomly position on a Si substrate[62]. In the used fabrication process, areal density but not pillar-pillar distance was controlled. Investigation of FAs showed that cells on the arrays with the intermediate areal density had the largest number of FAs that also had the most asymmetric shape. It was suggested that some of these FAs formed on the sidewalls of the nanocolumns. This was not observed for surfaces with low and high areal nanocolumn density.

In previous work we have described detailed protocols for fabrication of SU-8 polymer nanostructures on flat glass surfaces[63], and explored cell behaviour for two different cell lines on these surfaces[48, 45]. In this work, we use electron beam lithography (EBL) to fabricate surfaces decorated with vertically aligned SU-8 polymer structures to study changes in actin cytoskeletal and FA organisation in the osteosarcoma epithelial cell line U2OS. We perform both qualitative and quantitative analysis of the changes induced by the surface with different topological cues.

## 2 Materials and Methods

### Fabrication of nanostructures and sample mounting

SU-8 nanostructures were fabricated as previously explained[63]. Briefly, 24 mm by 24 mm glass cover slips (#1.5, Menzel-Gläser, thickness 170 μm) were cleaned by immersion in acetone, isopropyl alcohol, rinsed in de-ionised water and dried. The cover slips were then oxygen plasma treated for 2 min (Diener Femto plasma cleaner, power 100 W, base pressure 0.3 torr), followed by dehydration for 10 min on a 150 °C hot plate. Samples were then placed in a desiccator containing an open vial of Hexamethyldisilazane (HMDS, Sigma Aldrich product no: 440191). HMDS was applied by vapour deposition, the desiccator was pumped to low vacuum using a diaphragm pump for 5 min and the samples were kept in HMDS atmosphere for 60 min.

Substrates for EBL were prepared directly after HMDS treatment by spin coating SU-8 2001 (Microchem Corp.) to a desired thickness of 500 nm and 1000 nm. SU-8 was made fluorescent by adding either Oxazine 170 perchlorate, Rhodamine 800 or Coumarin 102 (all Sigma Aldrich) to a final concentration of 100 μg mL^−1^ resist. After spin coating samples were dehydrated on a hot plate at 95 °C. To mitigate charging during EBL exposure samples were then covered by a layer of conductive polymer AR-PC 5091 Electra 92 (AllResist GmbH) by spin coating at 2000 rpm for 60 s to thickness of 50 nm.

An Elionix ELS-G100 100 kV EBL-system was used to fabricate SU-8 nanopillars (NPs) with processing parameters as described in our previous work[63]. Table 1 summarise the arrays fabricated for this work. Pillar arrays were exposed using the Elionix dot-pattern generator where each pillar is exposed in a single exposure. Arrays were exposed over an area of 2000 μm by 4000 μm, with a current of 500 pA in write fields of 500 μm by 500 μm. NPs had a tip diameter of about 100 nm as a base diameter of 150 nm and 200 nm for structures of length 500 nm and 1000 nm respectively.

**Table 1:** Overview over pillar arrays used for the study of actin organisation and FA localisation. Arrays with a given NP height and array pitch are categorised into either Dense or Sparse and the corresponding density of NPs is reported.

| Height<br>[nm] | Pitch<br>[nm] | Category | Density<br>[NPs/$100 \mu\text{m}^2$] | Exposure time<br>[mm<sup>2</sup> / min] |
| --- | --- | --- | --- | --- |
| 500 | 500 | Dense | 456 | 0.7 |
|  | 1000 | Sparse | 115 | 1.5 |
| 1000 | 750 | Dense | 205 | 0.9 |
|  | 1000 | Dense | 115 | 1.5 |
|  | 2000 | Sparse | 29 | 4.0 |

After EBL exposure, the samples were rinsed in DI-water to remove the conductive polymer, then post exposure baked for 2.3 min at 95 °C and developed twice in mr-Dev 600 (Micro Resist Technology GmbH) developer for 20 s, rinsed in isopropyl alcohol and dried. Samples were then treated with oxygen plasma (Diener Femto plasma cleaner, power 50 W, base pressure 0.3 torr) for 30 s to render SU-8 hydrophilic and to give it similar surface chemistry as glass by oxidising surface epoxy-groups to hydroxyl.

Fabricated structures were imaged using Scanning electron microscopy (SEM) and samples sputter coated with 5 nm Platinum/Palladium alloy deposited with a 208 HR B sputter coater (Cressington Scientific Instruments UK). SEM was performed with a FEI Apreo SEM, at 5 kV and 0.2 nA with sample 45° pre-titled stage and with additional tilting of 30°.

When exposing the pillars, an indexing system was also exposed to make navigation during live-cell imaging more reliable. Arrays were optically inspected after fabrication to ensure free and standing pillars. The short Oxygen plasma treatment to render the SU-8 structures did not lead to any optically visible change to the structures. Lastly, the samples were mounted underneath 35 mm diameter dishes (Cellvis, Mountain View, CA, USA) with 14 mm holes and nano-structures pointing upwards, as indicated schematically in Figure 1. As flat surfaces, areas outside the structured part of the same samples were used. Before usage, all dishes were disinfected with 70% ethanol twice and dried.

**Figure 1:**
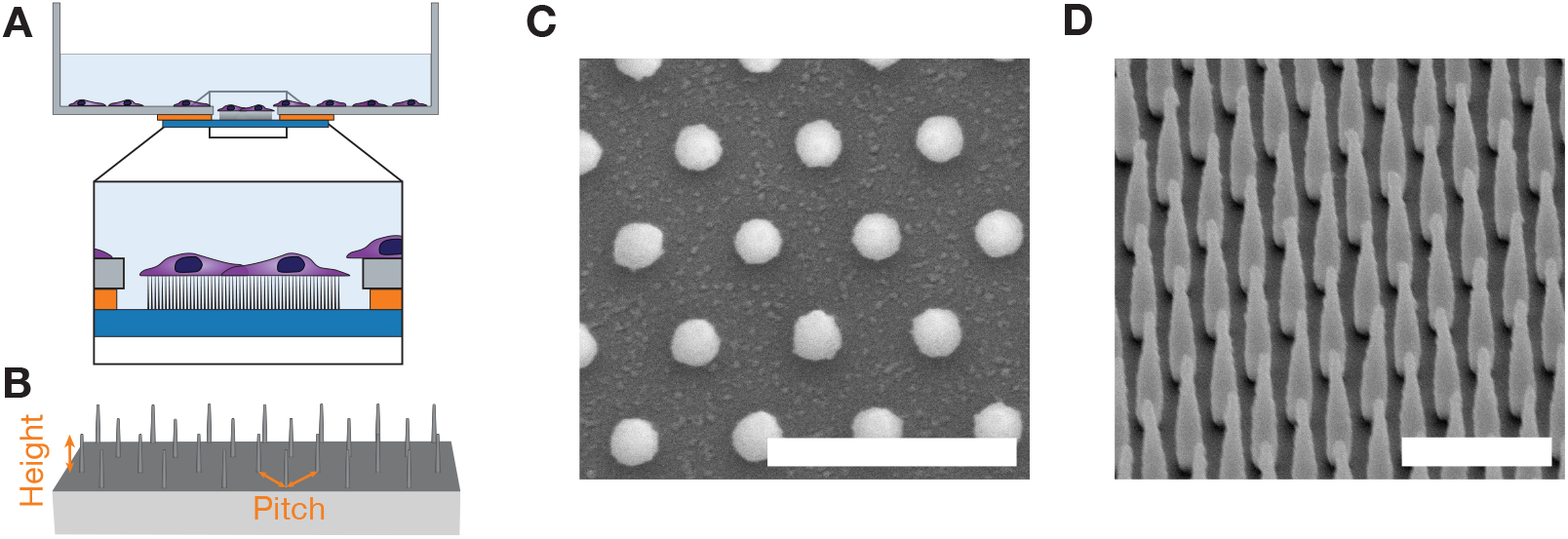
A) Side view schematic representation of nano-structured surface mounted in petri dish. Glass slides are mounted using paraffin such that structures are pointing upwards. B) Tilted schematic representation of nano-pillar array on flat surface, and two important parameters for the nano-pillar arrays (height and pitch). These figures are not drawn to scale. C-D) Overview of the nanopillar arrays employed in this work. Top-down and tilted side-view scanning electron micrographs of fabricated nano-pillar array with pillars of height 1000 nm and pitch 1000 nm. Scalebars 2000 nm.

### Cell culture and transfection

U2OS-cells (ATCC) were cultivated in Dulbecco’s modified Eagle’s Medium (DMEM Prod. 41965039, Fischer Scientific) with 10% fetal bovine serum (FBS) and kept at 5% CO_2_ and 37 °C. Before detachment, cells were washed with PBS and detached with Trypsin-ethylenediaminetetraacetic acid (trypsin-EDTA) and seeded on nanostrucutred or flat surfaces. For the diameter 14 mm glass wells 15 000 cells were seeded.

For the standard transfection experiments, cells were allowed 6 h for adhering to surfaces before transfection. U2OS cells were transiently transfected using Lipofectamine 2000 (Invitrogen, Fischer Scientific) by adapting the manufacturer protocol to our system. Briefly, 2 μL Lipofectamine 2000 was added to 50 μL Opti-MEM I Reduced Serum Media (Prod. 11058021, Gibco, Fischer Scientific) and incubated for 5 min at room temperature. Plasmid DNA coding for fluorescent LifeAct-TagGFP2 and TagRFP-vinculin fusion proteins were co-transfected by using 0.5 μg plasmid DNA (vinculin-pTagRFP and pCMVLifeAct plasmids) was diluted in 50 μL Opti-MEM I and incubated at room temperature for 5 min. For co-transfection of TagRFP-vinculin and pCMVLifeAct 0.5 μg of each plasmid was used.

The diluted DNA was added to the diluted Lipofectamine 2000 in a 1 : 1 ratio, and left to incubate for 20 min at room temperature. 40 μL of the combined transfection complex was then added to each well. After 18 h, 1.5 mL DMEM (Prod. 41965039) supplemented with 10% FBS and 1% 10 000 U/mL Penicillin-Streptomycin was added to each dish.

For reverse transfection experiments, the same amounts of reactants were used, but the transfection complex was added to a suspension of U2OS cells, and the suspension was then added to the wells.

### Microscopy

Live cell imaging was performed using a Zeiss LSM 800 Airyscan with an inverted Axio Observer Z1 stand connect to a PeCon compact incubator. Imaging was performed in an humidified environment at 37 °C, with 5% CO_2_ flow. High resolution imaging was performed using a Zeiss Plan-Apochromat 63x/1.4NA DIC M27 oil objective with Cargille Immersion Oil Type 37 (n = 1.51) suited for use at 37 °C. All images were taken using the system optimised pixel size both in-plane (typically 34 nm) and for stacks in the vertical axis (typically 180 nm).

To minimise imaging bias, imaging was performed in a standardised manner where each pillar array was raster scanned and cells expressing both LifeActGFP and Vinculin RFP were imaged. The high resolution images were then processed using a Zeiss algorithm for reconstruction of AiryScan images and exported as CZI-files for further manual and automatised image processing.

### Image analysis

For all cells, cell shape was based on the expression LifeActGFP fusion protein and expression of TagRFP-vinculin was used to identify FAs. Segmentation of images was performed using a script written in Python 3[64] using CZIfile[65] (version 2017.09.12) for reading the microscopy images in Zeiss-format. The python packages Scipy[66] and Scikit-image [67] were used for multi-dimensional image processing and image segmentation respectively.

To reduce the influence from fluorescence cross-talk from pillars (due to Oxazine 170 perchlorate, Rhodamine 800 or Coumarin 102), the pillar/surface channel was used as a background and subtracted from the TagRFP-vinculin imaging channel. A median filter (size: 10 pixels) was applied to remove noise from the TagRFP-vinculin channel, followed by classification of the image into regions based on their intensity value using a Multi-Otsu approach. Multi-Otsu thresholding with three classes was applied. The first class was typically the background, the second class constituted the cytosolic vinculin, whereas vinculin rich areas in FAs appeared brighter and could be classified into a third class. The quality of the image segmentation was briefly assessed by comparison to manual segmentation.

Area of cells and vinculin rich regions were described by counting pixel numbers and from this the actual area was found by correcting for the pixel size. Shape geometries were described by fitting each region with an ellipse with the same second-moment as the segmented region. In order to describe the cell area geometry, three measures were used: 1) Aspect ratio defined as the ratio of the ellipse major axis to the minor axis. 2) Circularity given as,

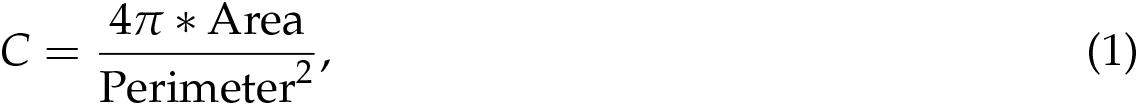

and roundness given as,

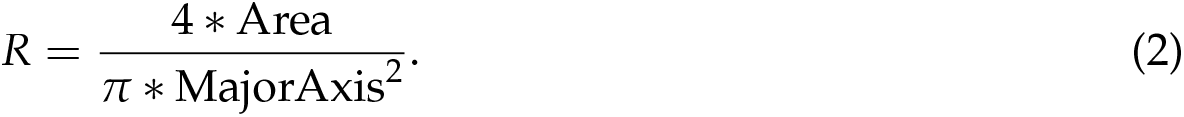

Segmented vinculin areas with a fitted ellipse that were too round (aspect ratio ≤ 1.5) or too elongated (aspect ratio ≥ 8.5) were rejected. In addition, vinculin areas smaller than 0.05 μm^2^ were filtered out. In order to find the distance between each vinculin area and the cell edge, the shortest euclidean distance between each centroid (the centre of the fitted ellipse for each vinculin area) and the cell edge was calculated.

### Statistical analysis

Statistical comparisons of distributions were performed by using the non-parametric two-tailed Mann-Whitney test neither assuming normal distribution nor equal standard deviation. P-values ≥ 0.05 were considered to represent a non-significant (ns) difference between the two populations. Significant values were denoted with * for *p* in 0.01 to 0.05, ** for *p* in 0.001 to 0.01, *** for *p* in 0.0001 to 0.001 and lastly **** for *p* ≤ 0.0001.

## 3 Results

Using previously established protocols, we have fabricated glass cover-slips decorated with precisely defined arrays of vertically oriented SU-8 nanopillars (NP) with variable separation and defined geometry[63]. Surfaces with NP areal densities of 456, 205, 115 and 29 NPs/100 μm^2^ (corresponding to pitches of 500 nm, 750 nm, 1000 nm and 2000 nm) were used. First we examine general trends in cell morphology, structure of actin cytoskeleton and cell-substrate interactions. We follow this by quantitative comparison of cell and FAs morphology on various nanostructured substrates and flat glass controls. We combine high resolution microscopy with high throughput fabrication to do qualitative analysis of at least ≈100 cells for each surface type, with imaging after 24 h and 48 h. In total we analyse > 400 high resolution images and 20 3D data sets.

Figure 1 shows a schematic representation of the NP arrays and electron microscopy images of fabricated substrates. Table 1 shows geometric parameters of arrays used in this work, their classification, as well as the corresponding NP area number density. We classify NP arrays into dense and sparse depending on observed cell adhesion behaviour (see below).

Figure 2 shows representative U2OS cells cultured on glass (A) and nanostructured surface (B-F) for 24 h. The cells have been co-transfected with pCMV-LifeAct-GFP and pTAG-RFP-Vinculin, that allows the visualisation of F-Actin and vinculin through production of flourescent LifeAct-TagGFP2 (hereafter: LifeActGFP) and TagRFP-vinculin fusion proteins respectively. F-actin network and vinculin rich areas in FAs are clearly detected. At the cell periphery, the LifeActGFP signal is in close proximity to the membrane and therefore we use this signal to visualise cell morphology. Signal from SU-8 NPs is shown in blue (see Experimental section).

**Figure 2:**
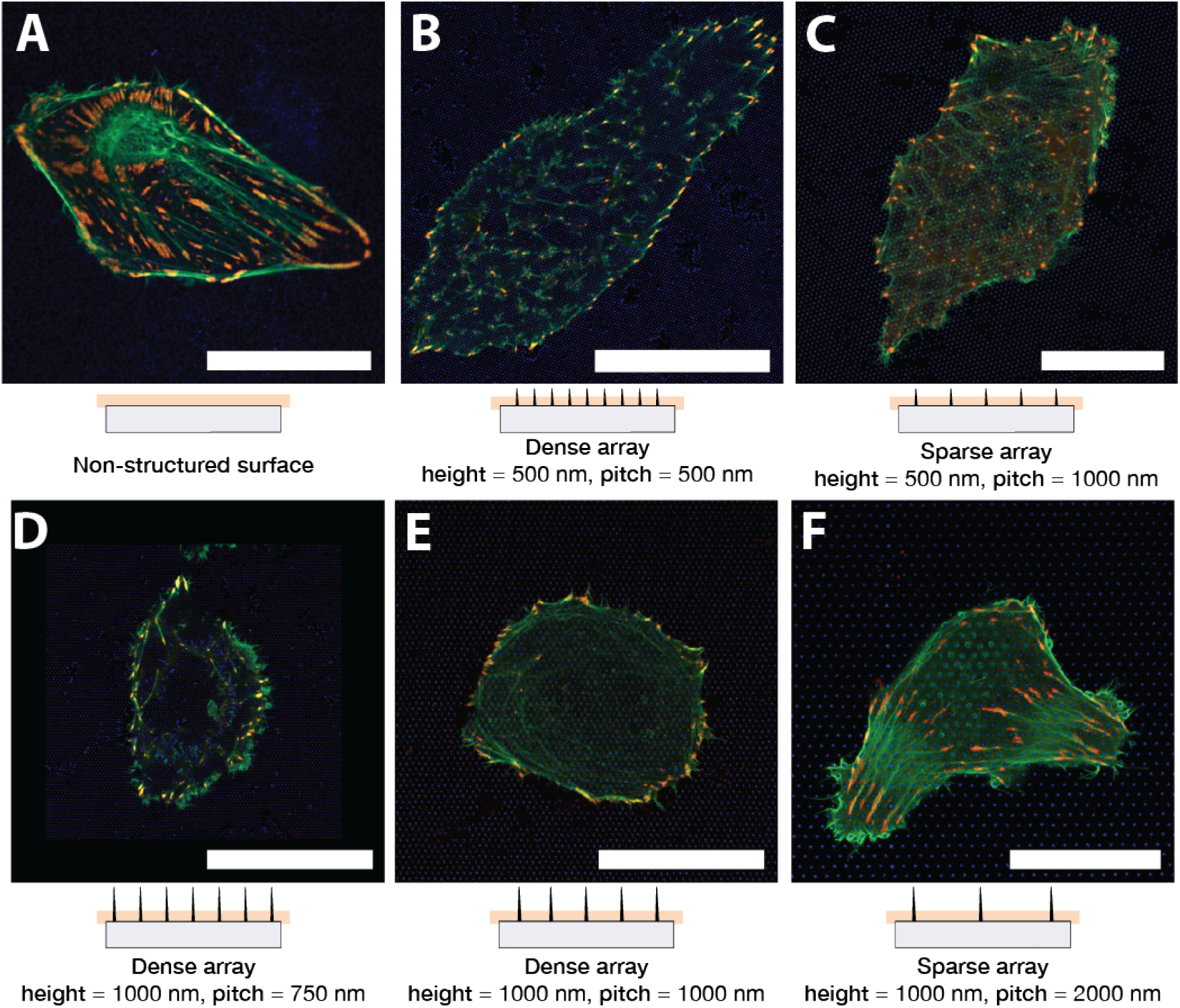
U2OS expressing fluorescent LifeActGFP (green) and TagRFP-vinculin (red) on different surface types. Yellow colouring indicates overlapping signals from LifeActGFP and TagRFP-vinculin channels. Below each micrograph a schematic side-view of the corresponding NP array is shown, together with the approximate position of the acquisition plane. Cells imaged on A) non-structured flat glass surface and on pillar arrays with different array pitch and structure heights: B) 500 nm pitch and 500 nm height, C) 1000 nm pitch and 500 nm height, D) 750 nm pitch and 1000 nm height, E) 1000 nm pitch and 1000 nm height, and F) 2000 nm pitch and 1000 nm height. All images presented are of representative cells. Scalebar 25 μm. Note that C) has a different scale than the other images.

Preliminary tests using transfected cells showed that cells seeded on both glass and structured surfaces appeared to be fully spread after approximately 6 h. In the following experiments cells were transfected 6 h after seeding, and then imaged 24 h and 48 h after transfection, corresponding to transfection 30 h and 54 h after seeding. In the following, these two time points will be referred by the observation time after seeding, that is 24 h and 48 h.

Cells were seeded on NP arrays with height 500 nm and 1000 nm and pitch 750 nm, 1000 nm and 2000 nm. After the initial spreading, cells were observed to be either round or elongated, similar to the situation on flat surfaces. This general morphology was found to be consistent over multiple experiments. Cells seeded on sparse NP arrays generally had a shape similar to the cells on glass surface, see Figure 2F depicting a representative cell on a 2000 nm pitched array. F-actin fibres were present also at the base of the NPs and in proximity of the glass surface, indicating that the cells were able to access the area close to the substrate. As observed previously[68, 62, 45], cells on dense arrays typically appeared to be suspended on top of the NPs (Figure 2B,Figure 2D and Figure 2E). Cells on dense arrays appear to have less prominent F-actin close to the glass surface, indicating that the actin fibres were not formed between pillars in proximity to the substrate. The relation between NP height and separation determined if the cells adhered to the substrate or were suspended on the top of the NP array. This is for example illustrated in Figure 2C where shorter NPs lead to the cell contacting the substrate, whereas longer NPs hindered contact, Figure 2E. These observations of actin fibres were further corroborated by z-stacks performed for some of the surfaces, as presented in Figure 3.

**Figure 3:**
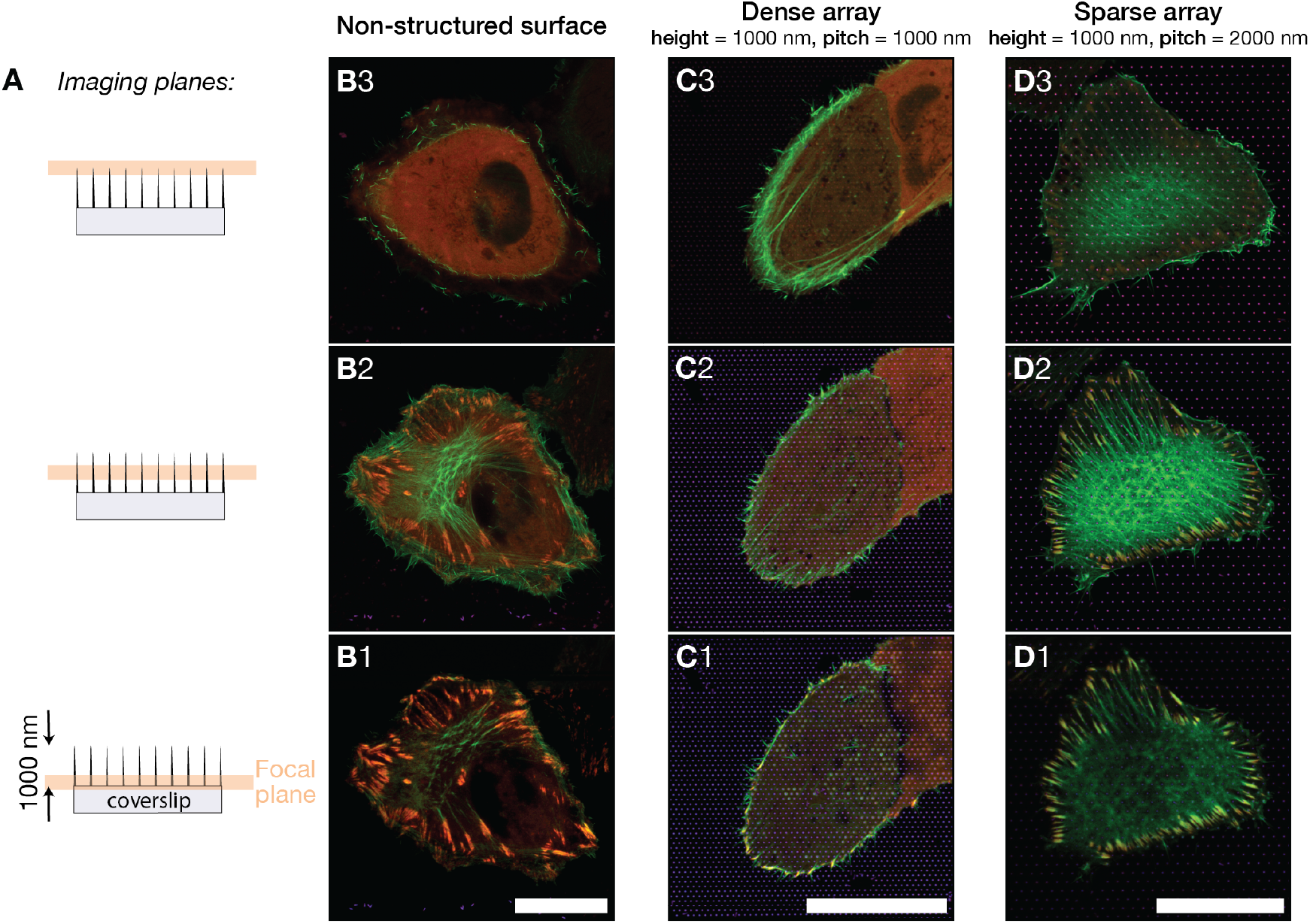
A) Images of U2OS cells on different surfaces were acquired at three indicated focal planes. Images were obtained close to the pillar base, approximately at half-pillar height, and close to the pillar apex. The imaging planes were located 0.0 μm, 0.4 μm and 0.8 μm from the flat glass surface. B,C,D) Merged fluorescence images showing U2OS expressing fluorescent LifeActGFP (green) and TagRFP-vinculin (red) on different surface types, yellow colour is indicative of overlapping LifeActGFP and TagRFP-vinculin. Imaged surfaces were B) non-structured glass surface, C) 1000 nm pitched pillar array, D) 2000 nm pitched pillar array. Presented images are representative for cells on each surface type. The vertical distance between focal planes for each surface type is approximately 400 nm. Scalebars 25 μm

To obtain a more detailed understanding on how cells adhere to the structured and non-structured surface, we evaluated distribution of FAs as visualised by the presence of TagRFP-vinculin fusion protein. Cells on flat surfaces typically formed elongated FAs distributed underneath the whole cell body, as shown in Figure 2A.

On sparse arrays, U2OS were able to contact the glass surface between NPs and adhered similarly to cells on glass, shown in Figure 2C and Figure 2F. For these NP arrays, FAs formed on glass in-between NPs, and the F-actin signal was also detected in the image acquired close to the base of the NPs. This indicated that the cells were able to bend the membrane around the nanostructures. Cells on denser arrays however, such as 750 nm and 1000 nm separation and 1000 nm height, were clearly hindered from adhering to the substrate between the nanostructurues, as shown in images Figure 2D and Figure 2E that was acquired close to the base of the NPs. However, around the periphery, the cells were typically able to attach to the substrate between the nanostructures forming FAs, often directed by the symmetry of the underlying pillar array.

Cells spreading on NP arrays with shorter length, and with an inter-pillar spacing of 1000 nm formed adhesions both towards the periphery and under the cell body. The F-actin fibre orientation was directed by the symmetry of the underlying array, as shown in Figure 2C. However, the location and orientation of vinculin containing FAs did not exhibit any clear pattern, with FA forming in-between NPs.

U2OS cells on 500 nm pillars with inter-pillar distance of 500 nm generally formed fewer and smaller adhesions compared to the planar surface as shown in Figure 2B. For cells seeded on this array, actin fibres were primarily observed in proximity to the glass surface at positions where they terminated in FAs. Again, this is a sign that the actin network was hindered from contacting the surface, and the cells were therefore assumed to be suspended on top of the array. However, cells appeared to have more intact F-actin network forming above the pillars. Parts of the actin network that was observed in-between the pillars, appeared to align with structures in the underlying NP array. This can be seen in Figure 2B, as F-actin fibres and FAs predominantly form along to one of the lattice direction, ie. parallel to open “lines”, of the pillar array.

On both dense and sparse arrays, we observed “ring-like” F-actin structures that formed around NP that protrudes upwards into the cell body. The F-actin ring structure appeared to be more prominent on sparse arrays, as shown in Figure 2E and Figure 2F. The formation of F-actin rings around NP has previously been described for fibroblasts on similar surfaces[45] and for U2OS cells on nanostructures with a range of structure sizes[58].

Based on the results presented above, we have selected three surfaces for a more detailed and quantitative description of cell morphology and FAs. We study cells on dense arrays (pitch 1000 nm, length 1000 nm), sparse arrays (pitch 2000 nm, length 1000 nm) and compare the results with cells on flat glass surfaces used as control.

By employing the Airyscan detector together with the dedicated image post-processing, we were able to perform imaging with an xy-resolution of about 140 nm and z-resolution of about 400 nm[69]. Figure 3 shows images of cells on the three surfaces, with imaging planes separated by approximately 400 nm. The investigation of F-actin bundles at different *z*. Cells on flat surface had a clearly visible F-actin network at the same focal plane or right above the FAs, (see Figure 3-B2). For cells on dense arrays, the F-actin network was found on a higher focal plane within the cell compared to the FA plane, which was in contact with the glass support (Figure 3-C1 and Figure 3-C2/C3). For cells on sparse arrays, the situation was similar to the cell on glass controls and actin network and FAs were detected at the same height (Figure 3-D2). This data support the initial observation that cells on sparse arrays attached the surface between the structures, whereas cells on dense arrays were primarily able to adhere to the surface around the cell periphery.

To analyse and quantify the differences in FAs and cell morphology for the three selected surfaces a Python based image analysis script was used (see Experimental). For the quantitative analysis, more than 300 high-resolution images were analysed. In these images, > 400 cells and > 7700 FAs were identified, Table 2 lists the number of detected cells and FAs for the three surface types included in the analysis. For all surfaces, cells were imaged both 24 h and 48 h after transfection. In the following analysis, geometrical parameters such as surface area, circularity and aspect ratio for cells on 3 surface types and after 24 h and 48 h are compared. Additional analysis can be found in the Supplementary Information. Surface area, circularity and aspect ratio for cells are shown in Figure 4 and for number of FAs, combined FA area per cell and fraction of FA area to cell in Figure 5. The geometrical parameters were defined as described in the Experimental section.

**Table 2:** Number of observations for cells (*n_cell_*) and for FAs (*n_FA_*) used in the quantitative analysis. Reported values correspond to the total number of cells identified by the image analysis script for each time point (24 h and 48 h) and surface type (flat surface, 1000 nm pillar array and 2000 nm pillar array).

| Surface | Time [h] | Observations |  |
| --- | --- | --- | --- |
| | | $n_{cell}$ | $n_{FA}$ |
| Flat | 24 | 29 | 1379 |
|  | 48 | 69 | 1280 |
| Pillars 2000 nm | 24 | 121 | 2513 |
|  | 48 | 94 | 1307 |
| Pillars 1000 nm | 24 | 80 | 814 |
|  | 48 | 30 | 451 |

**Figure 4:**
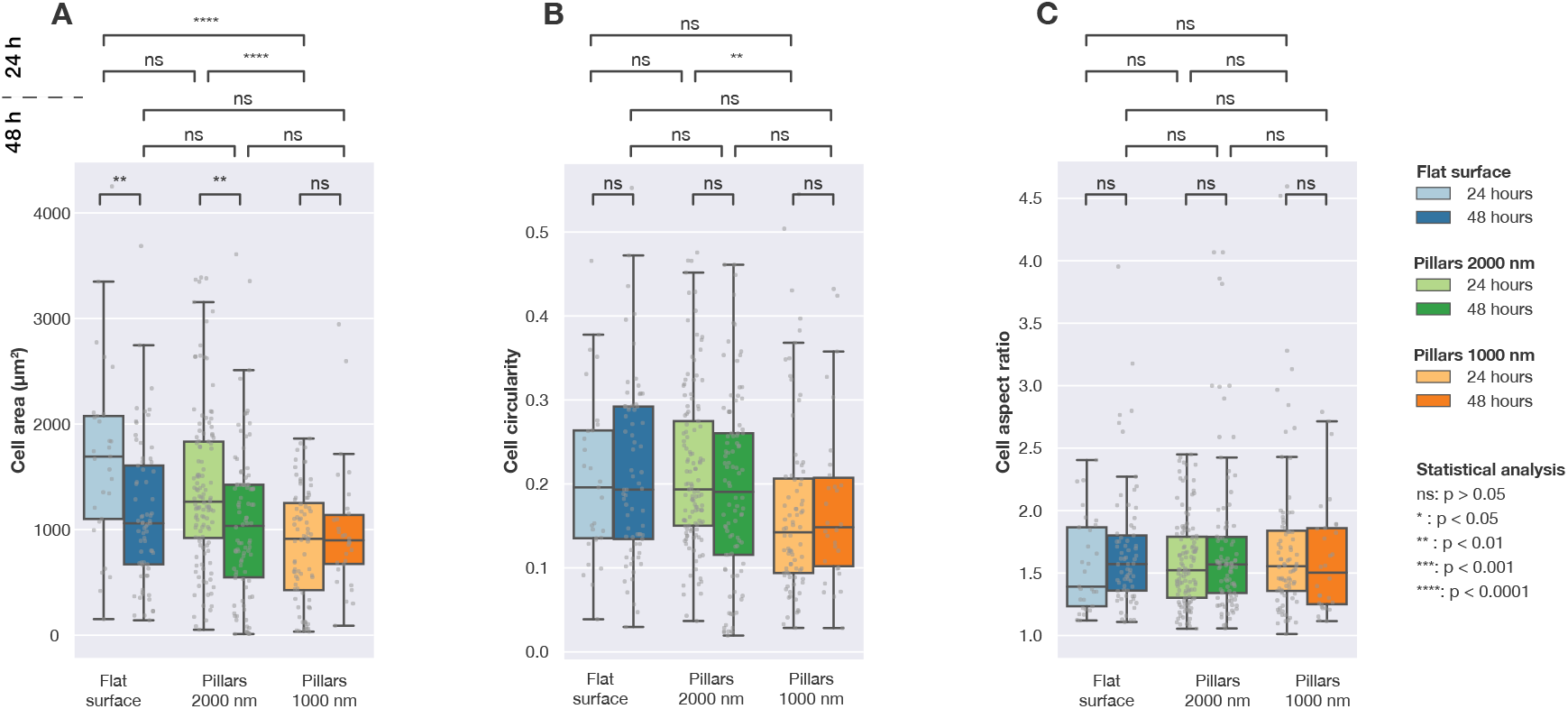
Calculated cell area, cell circularity and cell aspect ratio for U2OS cells expressing fluorescent LifeActGFP and TagRFP-vinculin fusion proteins on different surfaces imaged 24 h and 48 h post-transfection. The image detection was performed based on the intensity of the LifeActGFP signal. Each grey point correspond to one cell and the box plots shown median values (*Q*2) as well as the first (*Q*1) and third quartile (*Q*3). Statistical differences between the distributions were assessed using the Mann-Whitney non-parametric test.

**Figure 5:**
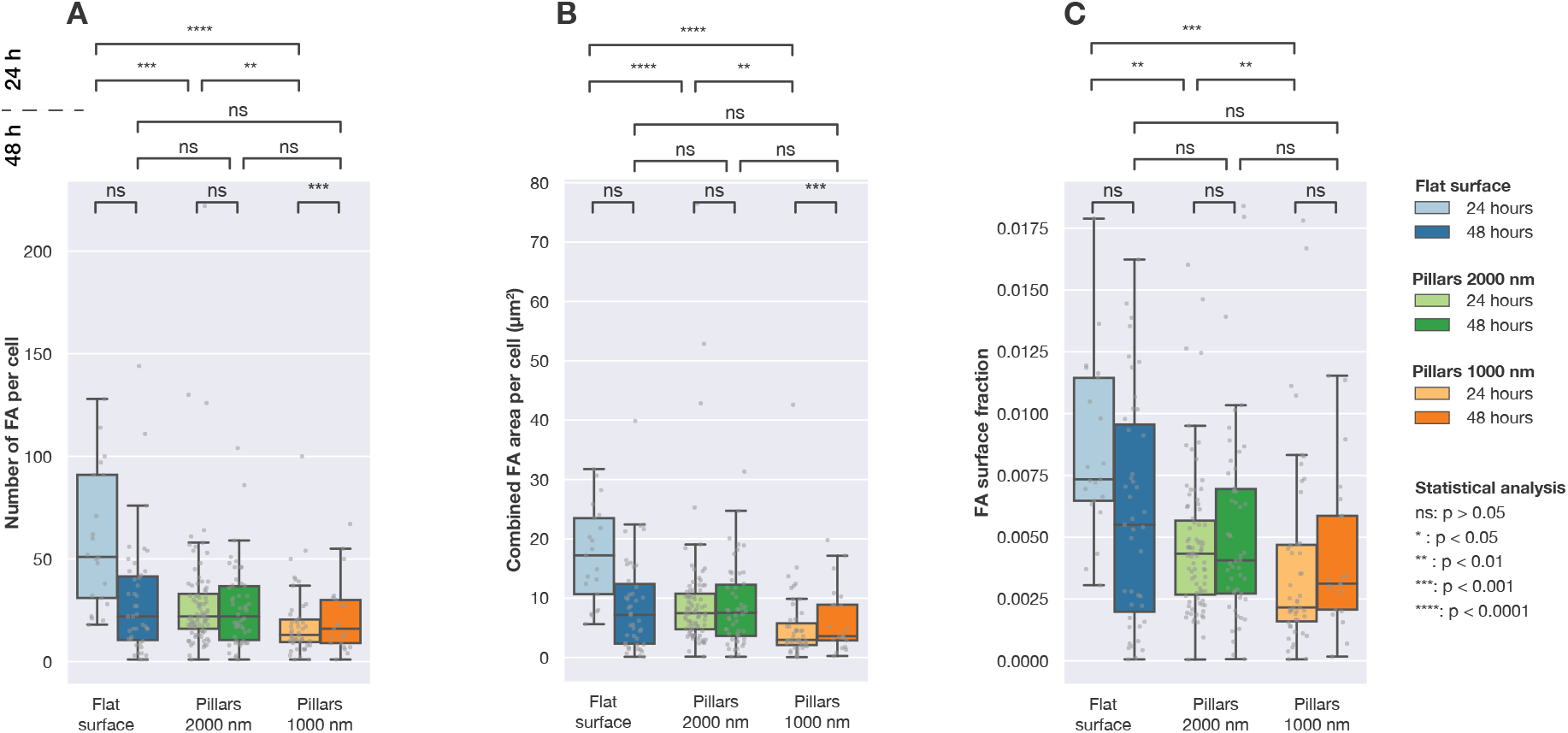
Number of FAs, combined FA area per cell and fraction of FA area to cell area for cells imaged on three different surface types (flat and pillar arrays with pitches of 1000 nm and 2000 nm). Each grey point corresponds to an observation from one cell. Statistical significance between the distributions was determined using Mann-Whitneys non-parametric test.

Figure 4 summarises data collected for both flat and structured surfaces. As shown in Figure 4A, significant differences in the cell area were observed after 24 h cell culture. However, after 48 h, there was no significant difference between average cell area on studied surfaces. When considering cell circularity (Figure 4B), no significant differences were detected between different surfaces, except between cells seeded on dense and sparse pillars imaged after 24 h. Cells on all three surfaces had the same average aspect ratios, as presented in Figure 4C.

Figure 5 shows the distribution of the number of detected FAs per cell, total surface area of FAs in each cell and the ratio of FA area to cell area. After 24 h the number of FAs formed by cells on the three different surfaces was significantly different. As shown in Figure 5B, the total FA surface area per cell was different for cells seeded on flat and structured surfaces. The same can be seen when comparing the relative amount of FAs (the total area of detected FAs divided by the total cell area) for the different surfaces, as shown in Figure 5C.

However, after 48 h of culture, significant differences between cell populations were no longer observed. When considering number of FAs per cell, combined FA area per cell or FA surface fraction, no differences between the three surfaces were found.

To understand whether the presence of NPs influence the localisation of FAs in the cell, we performed further analysis using the location of the FAs. Microscopy data indicated that FAs in cells on dense NP arrays were located closer to the cell periphery, as indicated by Figure 2 and Figure 3. To quantify this trend, we calculated the shortest distance from each FA to the cell edge. This was performed as illustrated in Figure 6. F-actin was used to determine location of the periphery and by constructing distance maps, the distance between each centre of a detected FA to the cell periphery was calculated. To account for differences in cell sizes, we normalised the distances between the detected cell edge and FA by the maximum distance from the edge to the geometric centre for each cell (a distance that is equivalent to radius for cells that have a circular shape). Data normalised by the maximum distance is presented in Figure 7 and Table 3. For cells on flat surfaces, the FAs are distributed more towards the centre of the cell, whereas on both on sparse 2000 nm arrays and dense 1000 nm arrays, the FAs are located closer to the cell periphery. This effect is most notable for dense arrays with 1000 nm pitch. Results of an alternative normalisation approach, where the FA locations were normalised by cell surface area, are included in Supplementary Information. This data shows he same qualitative trends as data shown in Figure 7.

**Figure 6:**
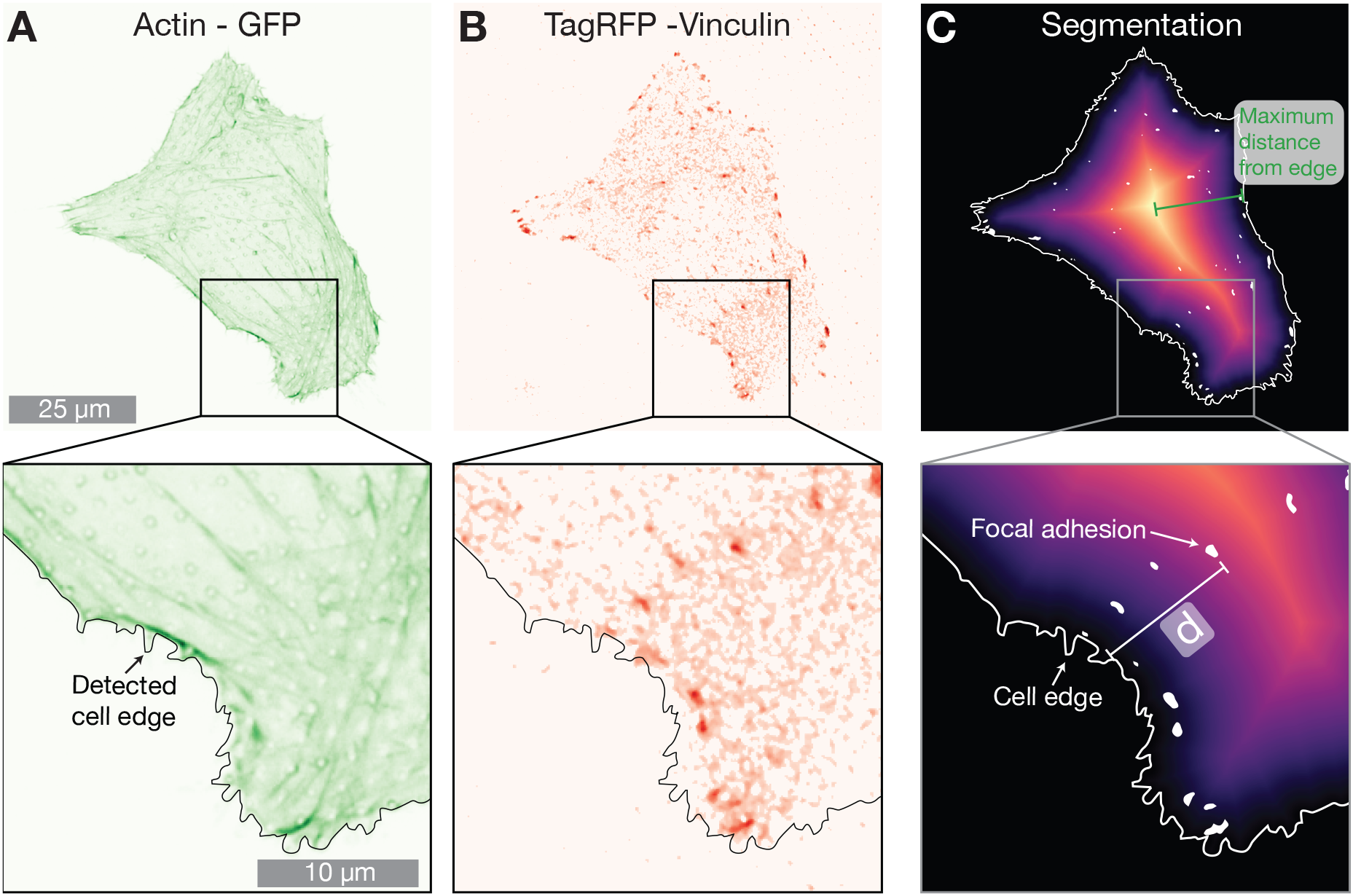
Example cell on 2000 nm pillar array expressing A) LifeActGFP (green) and B) TagRFP-vinculin (red). Superimposed on both figures is the detected cell edge determined by using the signal from expression of LifeActGFP (as shown in A). C) Shows a distance map from the detected cell edge as well as detected FAs. The shortest distance (denoted with *d* in figure) from each segmented vinculin spot (white areas) to the cell periphery (indicated with a solid white line) was calculated for all FAs in the images.

**Figure 7:**
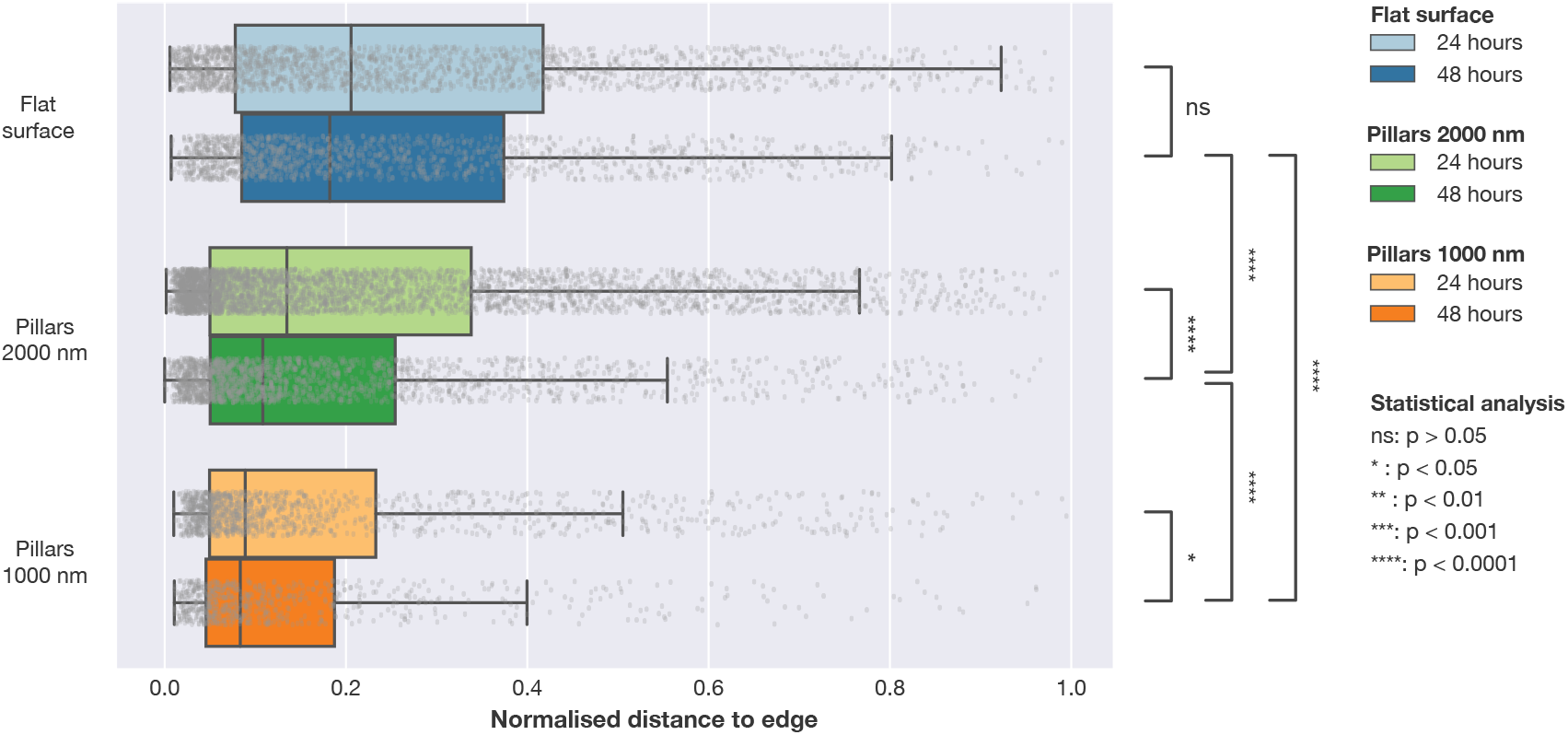
Distribution of FAs positions in relation to the closest cell edge normalised by the maximum distance from geometric centre of the cell to edge. Distances was obtained by calculating the distance from each observed FA to the cell edge defined by the LifeActGFP signal and plotted for the three surface types at 24 h and 48 h. Grey points represent individual observations of FAs and their distributions are summarised in the box plots. To test the likelihood that the FAs from the different surfaces and time points were from the same distribution, Mann-Whitney tests were performed on all distributions with significance levels denoted in the figure.

**Table 3:** Distance from the centre of a detected FA to the closest edge of cell defined by the LifeAct-GFP signal. Number of observations used in analysis is given as *n_FA_*. Values reported for the distance are median values (*Q*2) as well as the first (*Q*1) and third quartile (*Q*3).

| Surface | Time [h] | $n_{FA}$ | Distance [$\mu\text{m}^2$] | |
| --- | --- | --- | --- | --- |
|  |  |  | Q2 [Q1, Q3] |  |
| Flat | 24 | 1379 | 3.1 | [1.2, 7.7] |
|  | 48 | 1280 | 2.5 | [1.1, 5.8] |
| 2000 nm | 24 | 2513 | 1.9 | [0.8, 5.2] |
|  | 48 | 1307 | 1.4 | [0.8, 3.2] |
| 1000 nm | 24 | 814 | 0.9 | [0.6, 2.1] |
|  | 48 | 451 | 1.1 | [0.7, 2.1] |

## 4 Discussion

Organisation of actin cytoskeleton and formation of adhesions are processes studied extensively on flat surfaces. The present study was designed to investigate and quantify changes in organisation of the actin cytoskeleton and focal adhesions on nanopillar arrays.

Allowing the cells to spread and adhere to the surface for an extended period of time, makes it possible to observe the actin cytoskeleton organisation and the presence of fully matured FAs. 24 h after seeding, we observe significant differences in cell area, circularity and aspect ratio for cells seeded on the surfaces. However, no significant differences were detected after 48 h, indicating that the cells after 24 h have not yet fully adhered to the surfaces, and that the nanostructures mainly influences the FA organisation before they are fully matured. Neither after 24 h nor after 48 h is any FA formation on top or on the sides of the NPs observed.

Changes in the actin cytoskeleton organisation are also connected how a cell interacts with the surroundings. For example, both stress fibres and FAs grow when subject to stretching and appear to be functional interdependent[70]. Others have reported that rounder cells and FA localisation around edges are often observed for cells seeded on soft or compliant surfaces [71]. In our results, we observe a similar trend. Cells on dense arrays tend to show fibres around the cell edge, such as shown in Figure 2E or Figure 3B. FAs appear to form close to the cell edge for these cells. We speculate that when cells lack a flat surface, such as when the cells are suspended on top of the pillars, the FAs distribution resembles FAs on soft substrates, such as used by Prager-Khoutorsky and co-workers [71].

From our results, we observe that cells suspended on top of the pillars do appear to have a developed actin network above the pillars, but without FAs forming on the pillars themselves. For cells on sparse arrays however, cells appear to be less influenced by the NPs and both actin network and FAs appear more “flat-surface like”.

The interaction between FAs and actin cytoskeleton is complex and still not fully characterised. FAs linking the actin cytoskeleton to the ECM is known to act as traction points and to promote stress fibre formation in the cells. Conversely, actin fibres are again influencing the organisation and maturation of FAs. Numerous studies describe how cells tend to be suspended on top of dense NP arrays[62, 72, 61] and how cell membranes interact with single NPs[73, 74, 75]. These observations are corroborated by theoretical studies[68] and the cellular behaviour on pillars is fairly well understood. The mechanism behind FAs formation and attachment to the substrate around the cell edge on dense arrays remains unclear. In this respect, comparison with cells on a soft substrate is particularly interesting. For soft substrates, actin fibres are organised in a ring like fashion close to the cell edge and FAs form around the cell periphery[71]. On the nanopillar, arrays similar type of architecture is observed, but the actin fibres are typically shorter. Similar qualitative trends in terms of actin organisation and FA formation were observed by Li *et al.* for cells seeded on random nanowire arrays made from gallium phosphide [61].

Contrasting our results to other studies highlight an important aspect of studies on cellular response to NP arrays: cellular response may vary considerably depending on cell type, NP material, NP geometry and as well as other parameters. For example, Buch-Månson *et al.* studied fibroblasts and investigations of FAs showed that cells suspended on arrays with intermediate NP density had the highest number of FAs. In our results we do not see a similar trend. However, these studies cannot be directly compared as Buch-Månson *et al.* studied another cell line using a system with different array geometry, surface porosity and NPs length [62].

There are also studies describing the effect FAs placement has on cells[41]. By modelling cells on planar substrates Stolarska et al. suggest that the cells can control intra-cellular stresses by three mechanisms: FA position, FA size and attachment strength. FAs around the periphery allows the cells to be more sensitive to changes in the micro-environment. This could also be an underlying mechanisms for cells on NPs. Yet, it is not obvious that the results for the planar substrate are directly transferable to NP decorated surfaces.

Cell-interactions with the surrounding environment, for flat substrate, NPs arrays or *in vivo* ECM, are regulated by a complex set of relations between actin organisation, membrane mechanics, cell dynamics and contact with FAs. To further explore these relations, applying flat surfaces structured with NPs could be one promising approach. Such surfaces may also aid in exploring discrepancies in the cellular response to environmental cues between different cell lines.

## 5 Conclusion

In order to create more physiologically relevant systems for cellular studies, a plethora of 3D and 2.5D approaches have been proposed. One approach is to use flat-surfaces decorated with vertically aligned nanostructures as a simple model system. High resolution live cell imaging of co-transfected U2OS cells expressing pCMV-LifeAct-GFP and pTAGRFP-Vinculin have been used to study the influence of nanopillar arrays on actin cytoskeleton focal adhesion organisation. Our present results indicate that the U2OS cells spreading on surfaces decorated with nanopillars can be categorised into three different regimes by how they respond to the nano-structures. These observed changes are quantified by analysing more than 400 high-resolution images, and indicate that tuning geometrical properties of the nanostructured surface can be used to direct cell behaviour.

More specifically, the U2OS cells were found to either contact the substrate, attach preferably around the cell edge, or be fully suspended on top of the vertical NP arrays. In the latter case, we hypothesise that the resulting reorganisation of FA and cytoskeleton is an effect analogous to what is seen for softer substrates.

Increased understanding of how cells behave on nano-structured surfaces, such as pillar arrays, could help us discover more details about complex cellular processes. For example, it is still poorly understood how changes in the actin cytoskeleton and its architecture influence cell signalling. By studying the cell response on nanostructured surfaces in a systematic way, the potential connection between actin cytoskeleton, cell adhesions and a plethora of biochemical signalling pathways could be further explored. We therefore envision that further development of the presented platform and analysis could have implications for advanced *in vitro* applications or for development of smarter *in vivo* biointerfaces.

## Notes

### Competing Interest Statement

The authors have declared no competing interest.

