## Supplemental Information for "Analysis of actin and focal adhesion organisation in U2OS cells on polymer nanostructures"

### Supplementary Information: Actin cytoskeletal and focal adhesion organisation in U2OS cells on polymer nanostructures

2021-05-14

*

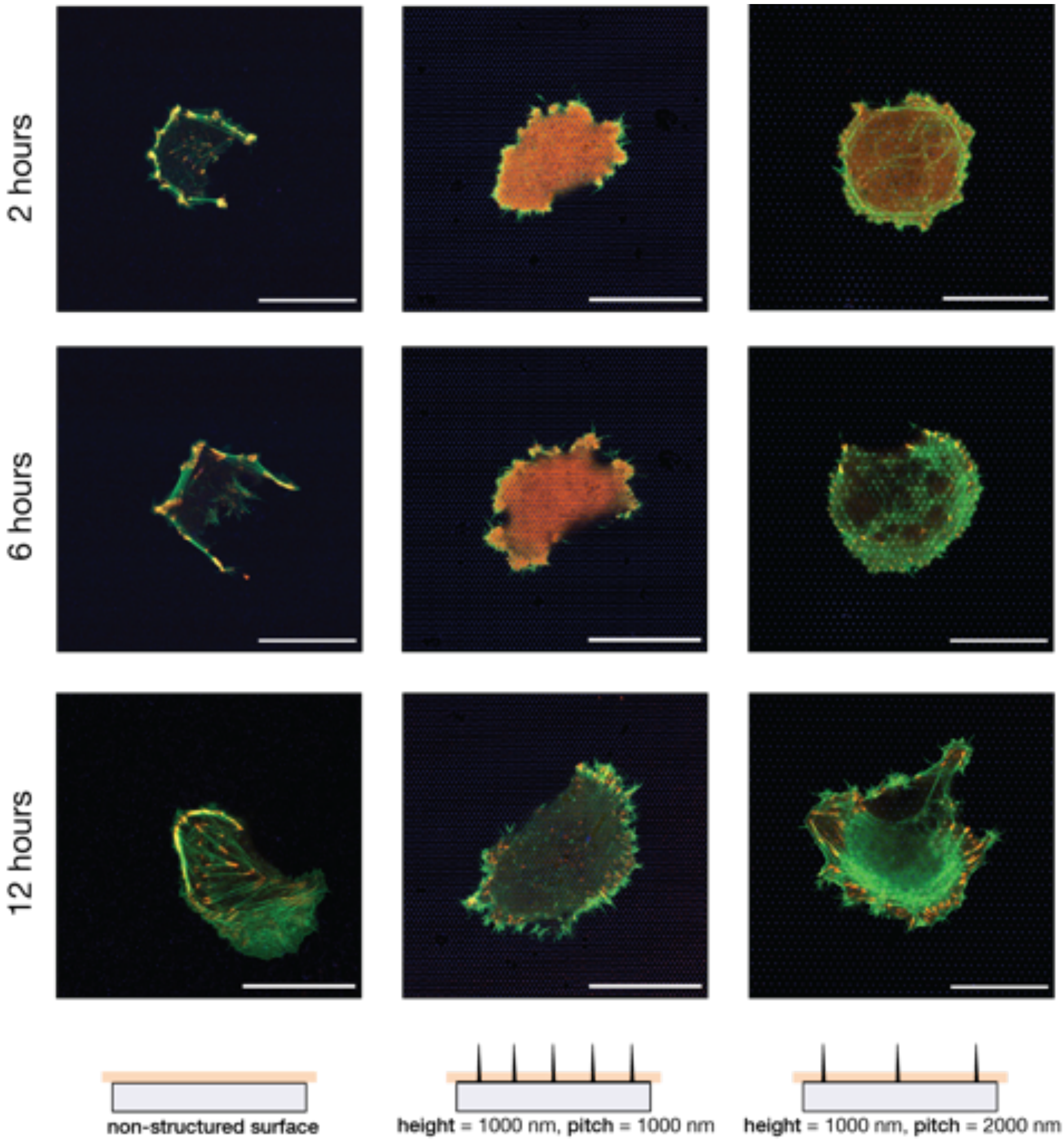


Figure S1: Cells transfected to express actin-GFP and TagRFP-vinculin fusion proteins imaged after 2 h, 6 h and 12 h. The same cell was imaged after 2 h, 6 h and 12 h. Scalebars 25 μm

Table S1: Number of detected cells for each combination of time after transfection and surface type segmented using the image analysis script. Reported values for cell area, circularity and aspect ratio are median values ($Q2$), with first ($Q1$) and third ($Q3$) quartile values in square brackets.

| Surface | Time [h] | Observations | Surface area [$\mu m^2$] | Circularity | Aspect ratio |
|---|---|---|---|---|---|
| | | | $Q2$ [$Q1, Q3$] | $Q2$ [$Q1, Q3$] | $Q2$ [$Q1, Q3$] |
| Flat | 24 | 29 | 1692 [1010, 2076] | 0.196 [0.135, 0.264] | 1.391 [1.234, 1.866] |
| | 48 | 69 | 1060 [671, 1607] | 0.193 [0.134, 0.292] | 1.572 [1.360, 1.801] |
| Pillars 2000 nm | 24 | 121 | 1264 [921, 1833] | 0.194 [0.150, 0.275] | 1.522 [1.302, 1.791] |
| | 48 | 94 | 1035 [548, 1425] | 0.191 [0.116, 0.260] | 1.569 [1.340, 1.789] |
| Pillars 1000 nm | 24 | 80 | 913 [427, 1252] | 0.142 [0.094, 0.206] | 1.555 [1.357, 1.839] |
| | 48 | 30 | 899 [675, 1139] | 0.148 [0.102, 0.207] | 1.503 [1.251, 1.859] |

Table S2: Summary of detected FAs. Number of observations corresponds to the total number of found FA for each combination of time point and surface type. Reported values for focal adhesion area, circularity and aspect ratio are median values ($Q2$), with first ($Q1$) and third ($Q3$) quartile values in square brackets.

| Surface | Time [h] | Observations | Surface area [$\mu m^2$] | Circularity | Aspect ratio |
|---|---|---|---|---|---|
| | | | $Q2$ [$Q1, Q3$] | $Q2$ [$Q1, Q3$] | $Q2$ [$Q1, Q3$] |
| Flat | 24 | 1379 | 0.231 [0.124, 0.425] | 0.598 [0.457, 0.761] | 2.314 [1.870, 3.036] |
| | 48 | 1280 | 0.218 [0.120, 0.435] | 0.618 [0.479, 0.772] | 2.144 [1.780, 2.781] |
| Pillars 2000 nm | 24 | 2513 | 0.260 [0.144, 0.448] | 0.650 [0.510, 0.804] | 2.174 [1.789, 2.832] |
| | 48 | 1307 | 0.253 [0.144, 0.442] | 0.638 [0.507, 0.792] | 2.142 [1.779, 2.722] |
| Pillars 1000 nm | 24 | 814 | 0.212 [0.134, 0.365] | 0.675 [0.517, 0.825] | 2.059 [1.622, 2.552] |
| | 48 | 451 | 0.242 [0.141, 0.421] | 0.669 [0.524, 0.820] | 2.069 [1.736, 2.597] |

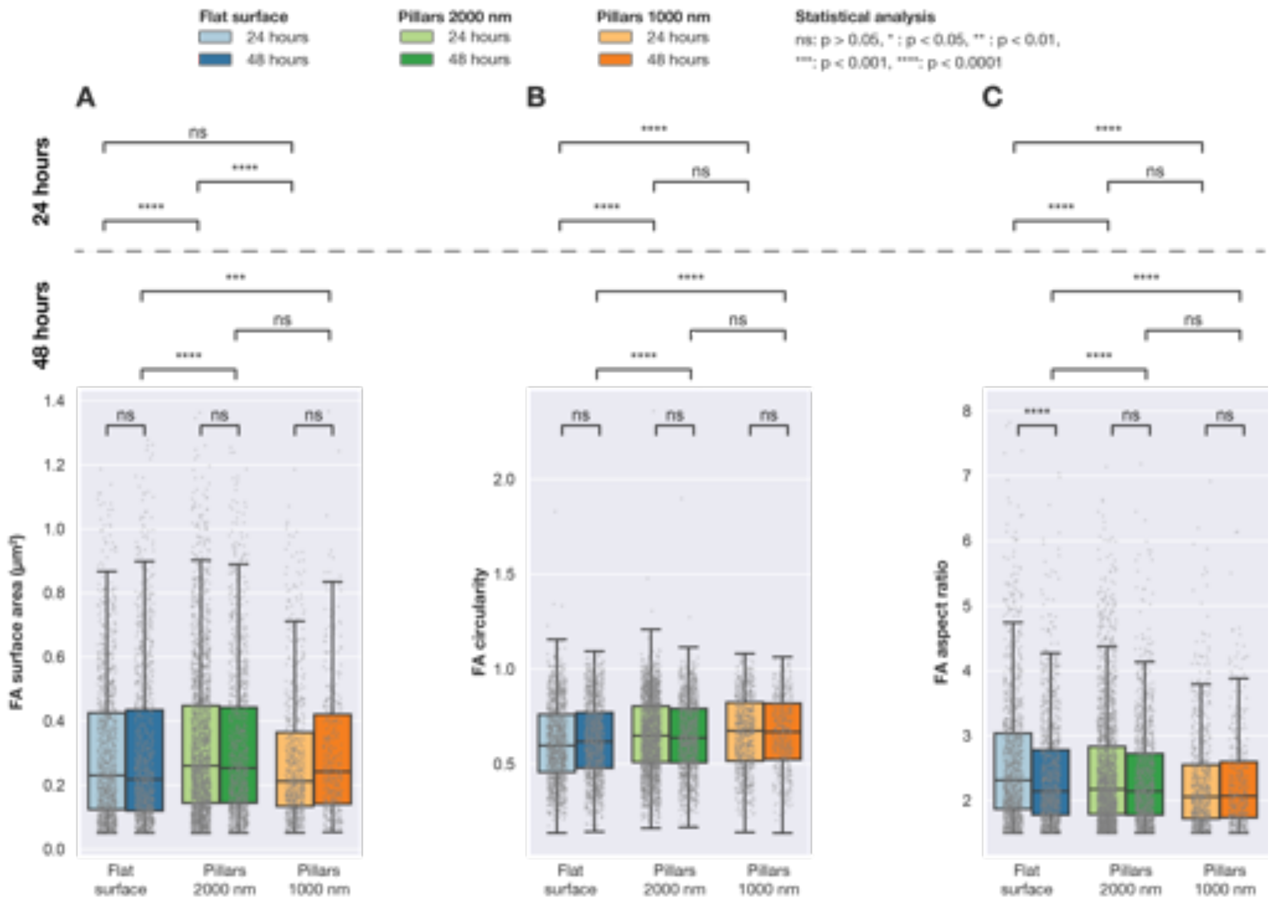


Figure S2: All detected adhesions detected in cells transfected to express fluorescent actin-GFP and TagRFP-vinculin fusion proteins were imaged after 24 h and 48 h. Based on the intensity of Vinculin, FAs were described by three geometrical properties: A) surface area, B) circularity and C) aspect ratio. Statistical differences between the distributions were assessed using the Mann-Whitney non-parametric test.

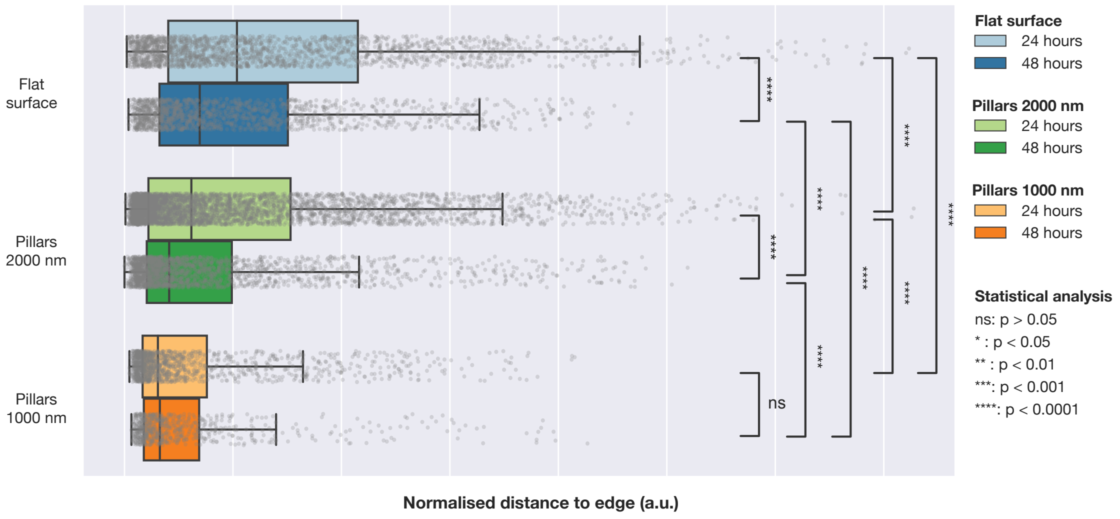


Figure S3: Distribution of FAs positions in relation to the closest cell edge normalised by cell surface area. Distances was measured by calculating the distance from each observed FA to the cell edge defined by the actin GFP signal and plotted for the three surface types at 24 h and 48 h. This distance was then normalised by the cell surface area for the specific cell. Grey points represent individual observations of FAs and their distributions are summarised in the box plots. To test the likelihood that the FAs from the different surfaces and time points were from the same distribution, Mann-Whitney tests were performed on all distributions with significance levels denoted in the figure.
